# Palmitic acid promotes glomerulosclerosis via TGF-β1 upregulation in podocytes

**DOI:** 10.64898/2026.11.14.766626

**Authors:** Rajesh Kavvuri, Adyasha Sarangi, Harshavardhan Komaragiri, Likhit A Reddy, Srividya Tamirisa, Anil K Pasupulati

## Abstract

Podocytes are terminally differentiated visceral epithelial cells of the glomerulus and play a vital role in renal filtration. Nevertheless, they are vulnerable to several insults in obesity and metabolic syndrome. Obesity-related glomerulopathy (ORG) is characterized by podocyte hypertrophy, reduced podocyte density, and secondary focal segmental glomerulosclerosis (FSGS). However, how lipid overload, particularly palmitic acid (PA), whose plasma levels are frequently elevated in obesity, injures podocytes remains incompletely understood. Human podocytes were exposed to PA (100 µM for 24 hours), and we assessed lipid accumulation, WT1 expression and promoter activity, TGF-β1 signaling, and cell-cycle regulators. PA induced lipid droplet accumulation and increased WT1 expression in podocytes. Concurrently, PA treated podocytes showed cytoskeletal aberrations, loss of podocyte structural genes (podocin, nephrin and synaptopodin) and TGF-β1/Smad2 axis causes upregulation of fibronectin, collagen, and α-SMA. Similarly, mice fed a high-fat diet (HFD) for 16 weeks exhibited TGF- β1 activation in podocyte enriched glomeruli together with glomerulosclerosis, impaired kidney function, and podocyte loss. In both models, Cyclin D1, Cyclin B1, CDK4, p53, and p21 were increased. Notably, ∼30% of PA-treated podocytes were binucleated, yet podocyte number did not increase. Lipotoxic stress was associated with concurrent induction of WT1 and TGF-β1, loss of the podocyte phenotype, and abortive cell-cycle re-entry accompanied by binucleation and podocyte loss. These findings point to a WT1–TGF-β1–cell-cycle axis as a potential contributor to ORG and a candidate therapeutic target. Whether WT1 directly regulates TGF-β1 remains to be determined.

## Introduction

As the third leading cause of death worldwide, kidney diseases imposes a substantial health and economic burden [1]. Obesity is a major and growing risk factor for chronic kidney disease (CKD), and it is associated with a distinct glomerular lesion known as obesity-related glomerulopathy (ORG), characterized by glomerulosclerosis, glomerulomegaly, podocyte injury, and proteinuria [2]. Obesity is accompanied by dyslipidemia and elevated circulating free fatty acids (FFA), particularly saturated fatty acids. Palmitic acid (PA) is the most abundant FFA, and its plasma levels are significantly elevated in obese patients [3]. PA is the first saturated fatty acid synthesized de novo; it also serves as a precursor for synthesis of several long-chain fatty acids. It is abundantly found in several commonly consumed foods such as butter, palm oil, and animal fat. Excessive dietary intake of PA-rich foods may adversely affect renal health.

Excess PA is not merely a metabolic fuel overload; it is a potent lipotoxic agent that drives oxidative stress, inflammation, and death of multiple renal cell types. PA acts as a TLR4 agonist, activating the TLR4/JNK/NF-κB signaling cascade and promoting the production of inflammatory mediators, including IL-6, a key contributor to inflammation [4], [5]. Elevated circulating PA levels are significantly associated with obesity, metabolic syndrome, and insulin resistance [6]. Intraperitoneal PA administered to female rats resulted in lipid dystrophy, oxidative stress, renal inflammation, and electrolyte imbalance [4]. PA was shown to exacerbate proteinuria-induced cell death and inflammation via the CD36-inflammasome axis in proximal tubular cells of obese mice [7]. PA selectively causes cell death in kidney podocytes by generating mitochondrial reactive oxygen species, reducing ATP synthesis, causing endoplasmic reticulum stress, and triggering the caspase-dependent mitochondrial apoptotic pathway [8]. Excessive PA led to local insulin resistance, loss of slit diaphragm proteins (nephrin and podocin), and disorganization of the actin cytoskeleton [9]. Either the depletion of podocytes or the structural collapse of podocytes induced by PA-mediated lipotoxicity could prevent the kidney from effectively retaining proteins, ultimately contributing to kidney dysfunction. Understanding insights into podocyte injury in the setting of ORG is of intense interest owing to the magnitude of obesity and CKD.

In an earlier study using spontaneously obese WNIN/Ob rats and high-fat-fed mice, it was found that ORG is associated with podocyte foot process effacement and lower expression of podocyte markers. Both models also showed raised expression of WT1, a transcription factor that is normally minimal in adult podocytes [10]. Nevertheless, in the above study, the specific effect of PA on podocytes was not investigated despite being the fact, the serum PA levels are significantly high among obese patients and associated with poor renal outcome [9], [11]. Podocytes are terminally differentiated, post-mitotic (quiescent) epithelial cells of the glomerulus. Podocytes, along with the glomerular basement membrane and capillary endothelial lining, form the final barrier that prevents protein loss through their interdigitating foot processes and slit diaphragms. Because podocytes cannot readily regenerate, their loss or detachment leads directly to proteinuria and progressive glomerulosclerosis. Maintenance of the quiescent state depends on cyclin-dependent kinase inhibitors (such as p27 and p57) that restrain entry into the cell cycle [12]. Cell cycle reactivation is therefore increasingly viewed as a maladaptive response contributing to podocyte depletion in several glomerular diseases. TGF-β1 is a central profibrotic cytokine in the kidney and a well-established mediator of podocyte injury. In podocytes, TGF-β1 signaling induces cytoskeletal rearrangement, apoptosis, loss of slit diaphragm proteins, and detachment, and it is upregulated in diabetic and other proteinuric kidney diseases [12], [13]. TGF-β1 has also been shown to modulate cell- cycle regulators, raising the possibility that it links metabolic stress to aberrant cell-cycle activity in podocytes [14]. Whether lipotoxic stress from PA engages the TGF-β1 pathway in podocytes, and whether this converges on cell cycle reactivation as a mechanism of injury, remains to be defined.

Here, we hypothesized that excess PA upregulates TGF-β1 in podocytes, thereby triggering cell cycle reactivation and subsequent podocyte injury. To test this, we exposed cultured podocytes to PA and examined TGF-β1 expression and signaling, cell cycle regulators, and markers of podocyte damage. We then fed mice a saturated fatty acid enriched diet and assessed the development of proteinuria and podocyte injury i*n vivo*. Our findings identify a TGF-β-dependent, cell-cycle-driven mechanism by which excess saturated fatty acids damage podocytes, providing mechanistic insight into ORG.

## Materials and methods

### Materials

The primary antibodies are as follows: The anti-p53 (ab26), anti-cyclinB1 (ab72), anti-p21 (ab109520), anti-CDK4 (ab137675), and anti-cyclinD1 (ab16663), were purchased from Abcam (Cambridge, MA). Anti-α-SMA (#48938), anti-SMAD4 (#46535), and anti-β-actin (#4970) were purchased from Cell Signaling Technology (Danvers, MA). The anti-Nephrin (NBP1-77303), and anti-podocin (NBP2-75624) from Novus Biologicals. The anti-TGF-β1 (MAB240) from R&D Systems (Minneapolis, MN). The anti-TGF-β1 for immunoblotting and immunohistochemistry (MA5-16949). Mouse/Rabbit Poly-Detector DAB HRP Brown Immunochemistry detection system (BSB020, Santa Barbara, CA). Phalloidin fluorescein isothiocyanate labeled (P5282) and glutaraldehyde solution (G5882) were obtained from Sigma Aldrich (St. Louis, MO, USA). Vectashield mounting medium containing 4′, 6- diamidino-2-phenylindole (DAPI) as a nuclear counterstain was procured from Vector Laboratories (Burlingame, CA). DyLight 488 and DyLight 564, and Cy5-conjugated secondary antibody were purchased from Vector Laboratories (Burlingame, CA). Paraformaldehyde (P6148-500G), bovine serum albumin (BSA, A3294-50G), fluorescent phalloidin–TRITC conjugate (P1951), 4, 6-diamidino-2- phenylindole (DAPI, P36971). All other reagents used were of analytical grade and obtained from Sigma Aldrich (St. Louis, MO, USA). Protease inhibitor cocktail was purchased from Roche (#11873580001, Basel, Switzerland). Urinary albumin (#11573) and creatinine (#11502) detection kits were procured from Biosystems (Barcelona, Spain). Avidin and biotin (#927301) and anti-mouse IgG (#406604) purchased from BioLegend (San Diego, CA). High-Fat Diet was purchased from Research Diets (#D12108C, New Brunswick, USA). Precision Plus Protein Dual Color Standards (Bio- Rad, Hercules, CA). BCA reagent (#786-570) and chemiluminescence substrate (#786- 081) were procured from G-Biosciences, (Geno Technology Inc., USA). Nitrocellulose membrane was purchased from Bio-Rad (#1620112, Germany). Trizol was purchased from Takara (#9108-RNAiso PLUS, Japan).

### Animal experiments

Male C57BL/6 mice (3 weeks old, weighing 14 ±3 g) were procured for diet-induced obesity studies and acclimatized for one week in the animal house facility at the University of Hyderabad. The animals were maintained under controlled environmental conditions, with a temperature of 22 ±2^0^C, relative humidity of 50%, and a 12h light/dark cycle. Following acclimatization, mice were fed a high-fat diet (HFD; 40% high-energy diet; Research Diets, #D12108C, New Brunswick, USA). Urine samples were collected after 12-16 weeks of HFD feeding. Blood samples were collected from the tail vein, and plasma was subsequently isolated. Mice (n=8 per group) were euthanized by carbon dioxide (CO_2_) asphyxiation, followed by perfusion prior to kidney harvesting. For histological analysis, kidney tissues were fixed in 4% paraformaldehyde, embedded in paraffin, and sectioned transversely at a thickness of 4 µm. The sections were mounted onto silane-coated slides. All animal experiments were conducted in accordance with the guidelines approved by the Institutional Animal Ethics Committee (IAEC), University of Hyderabad [15].

### Estimation of biochemical parameters

The urinary albumin and creatinine, estimated using a commercially available kit (#11573 & #11502, Biosystems) and reported as µg/mL.

### Podocyte cell culture and experimentation

In this study, conditionally immortalized human podocyte (a gift from Prof. Moin Saleem, University of Bristol) cells were cultured in RPMI 1640 containing 10% FBS and antibiotic & antimycotics under 5% CO_2_. Cells were maintained at 33^0^ C a permissive condition for proliferation and shifted to non- permissive 37^0^ C for 10-14 days to differentiate into mature podocytes. Differentiated podocyte cells were treated with or without Palmitic acid for 24 hours and harvested later for further analysis. For reporter assay and over expression analysis, transfection studies were performed in podocytes after 10 days of differentiation according to the manufacturer protocol using Lipofectamine 3000 from Invitrogen (#L3000008).

### Palmitic acid preparation

75mM Sodium palmitate (#P9767, sigma) was prepared by dissolving it in milli Q water at 72^0^ C as a stock solution and maintained until the solution becomes clear. Similarly 3.8% free fatty acid BSA (#HY-D0842A; MCE) prepared by dissolving in milli Q water at 40^0^ C. Further 4mL of 75 mM sodium palmitate transferred into 100 mL of 3.8% FFA BSA for conjugation without changing any temperature, to get a 3 mM working stock. The conjugated FFA BSA:PA were incubated in shaker incubator for 3 hours at 37^0^ C and 180 RPM. The conjugated solution was sterile filtered and stored in -20^0^ C. For the treatment, 100 uM of FFA BSA:PA was used as an inducer whereas 3.8% FFA BSA as a vehicle.

### Histological Examination and immunostaining

Paraffin-embedded kidney sections were subjected to hematoxylin and eosin (H&E) and Masson’s trichrome staining, as described previously [15]. Periodic acid-Schiff (PAS) was also performed according to previously established protocols. Following staining, the sections were dehydrated through graded alcohols, cleared in xylene and mounted using DPX mounting medium. For immunohistochemical detection of proteins, kidney sections were deparaffinized, rehydrated, and treated with 10% hydrogen peroxide in methanol for 20 min to block endogenous peroxidase activity. Antigen retrieval was performed using pre-boiled citrate buffer, followed by thorough washing and treatment with avidin and biotin. The sections were incubated overnight with respective primary antibodies diluted in 1% BSA followed by secondary antibodies for 1h after proper washes. Immunoreactivity was visualized using 3’3- diaminobenzidine (DAB) as the chromogenic substrate. The stained sections were examined using an EVOS M5000 microscope (Invitrogen, Carlsbad, CA, USA), and captured at 400x magnification.

### Immunofluorescence

Paraffin-embedded tissue sections were deparaffinized, rehydrated and subjected to antigen retrieval, similarly cells were grown on coverslips, followed by treatment as mentioned above and subsequent fixation with paraformaldehyde (4%). Both the sections and cells were blocked with PBS containing BSA (3%) before overnight incubation with primary antibodies. The next day, for F-actin staining the podocyte cells were incubated with fluorescent phalloidin–FITC along with Alexa Fluor conjugated secondary antibodies and tissues were alone with secondary antibodies for 1 h at room temperature. Next counterstaining with DAPI, followed by mounting with anti-fade mounting media. Fluorescence images were acquired using a Leica fluorescence microscope (Leica Microsystems, Wetzlar, Germany) at 400x magnification. Images were analysed using ImageJ (NIH).

### Isolation of Kidney Glomeruli

Glomeruli were isolated from kidney tissues using a series of stainless-steel sieves, as described previously [15]. Briefly, decapsulated kidney slices were placed on a 200 µm sieve and gently pressed using a sterile pestle, followed by washing with Hank’s balanced salt solution (HBSS). The resulting filtrate was subsequently passed through a 150 µm sieve. Glomeruli retained on the 75 µm sieves were collected as a suspension and allowed to settle for 10 min. The suspension was then centrifuged at 1000 rpm for 5 min, and the resulting pellet was collected for subsequent immunoblotting analysis.

### Cloning of WT1 promoter and gene

WT1 promoter (1264 bp, ranging from 0 to -1263) was cloned into the pGL3 basic vector and transformed into DH5-α cells. It was confirmed by the sequencing and used for reporter assay analysis. WT1 (+KTS isoform-927bp) was used for overexpression studies and prepared as mentioned previously [15].

### Immunoblotting

Extracted glomeruli from mice and cultured podocytes were lysed with RIPA buffer and protein was estimated using bicinchoninic acid method. Following protein quantification and normalization, equal amounts of protein from each sample were separated by SDS-PAGE. The resolved proteins were subsequently transferred into nitrocellulose membrane by electroblotting at 50V for 1.5h at 4°C. Blocking was done with 5% skimmed milk and incubated with respective primary antibodies for overnight at 4°C, followed by secondary incubation for 1h at room temperature. Bands were detected using a chemiluminescence substrate and visualized by ChemiDoc XRS System (Bio-Rad). Densitometric analysis was done using Image J, NIH. To assess the extent of proteinuria, equal volumes of urine samples were subjected to SDS-PAGE, followed by silver nitrate staining to visualize urinary protein profiles.

### RNA-isolation and qRT-PCR

Total RNA was isolated from immortalised podocyte cells with or without transfected with WT1 plasmid by phenol-chloroform method and expression of specific genes were analyzed by the qRT-PCR (QuantStudio from Applied Biosystems). Primers include WT1 (FP: CGAGAGCGATAACCACACAACG and RP: GTCTCAGATGCCGACCGTACAA) ACTB (FP: CACCATTGGCAATGAGCGGTTC and RP: AGGTCTTTGCGGATGTCCACGT), N-cadherin (FP: TGGAAATCAAGTGGAGAACCC and RP: TATGTTCCAGGCTTTGATCCC) COL1A1 (FP: TGAAGGGACACAGAGGTTTCAG and RP: GTAGCACCATCATTTCCACGA), COL4A1 (FP:AGCGAGATGTTCAAGAAGCC and RP: TGTTCTTCTCATACAGACTTGGC),Fibronectin (FP: ACAACACCGAGGTGACTGAGAC and RP: GGACACAACGATGCTTCCTGAG) and TGF-Beta1 (FP: CATCAACGGGTTCACTACCG and RP: CCATGAGAAGCAGGAAAGGC).

### Statistical analysis

Data are presented as Mean±SD. GraphPad Prism 7 software was used to analyze statistical differences between the distribution of two (Unpaired nonparametric-Mann Whitney test) or multiple independent samples; we used one-way ANOVA (Tukey’s Multiple comparison test). p ≤ 0.05 was considered significant

## Results

### PA treated podocytes showed lipid accumulation and elevated WT1 expression

Podocytes exposed to PA (100 µM for 24 hours) showed ectopic lipid accumulation (lipotoxicity) in podocytes as evidenced by Oil Red O staining, which is further confirmed by BODIPY staining (Figure 1A&B). Ectopic lipid accumulation in PA exposed podocytes showed elevated levels of WT1, but decreased expression of podocyte specific markers such as nephrin, podocin, and synaptopodin (Figure 1C&D). Besides elevated WT1 expression, the immunoflourescence analysis revealed increased podocyte cell size upon exposure to PA as evidenced by increased nucleus to cytoplasmic ratio (Figure 1E&F). Phalloidin staining revealed disruption of actin stress fibers in PA treated podocytes (Figure 1E). It is also observed that in approximately 30% of cells that were treated with PA showed binucleated podocytes (Figure 1E). We also observed that increased WT1 promoter activity in podocytes exposed to PA (Figure 1G).

**Figure 1:**
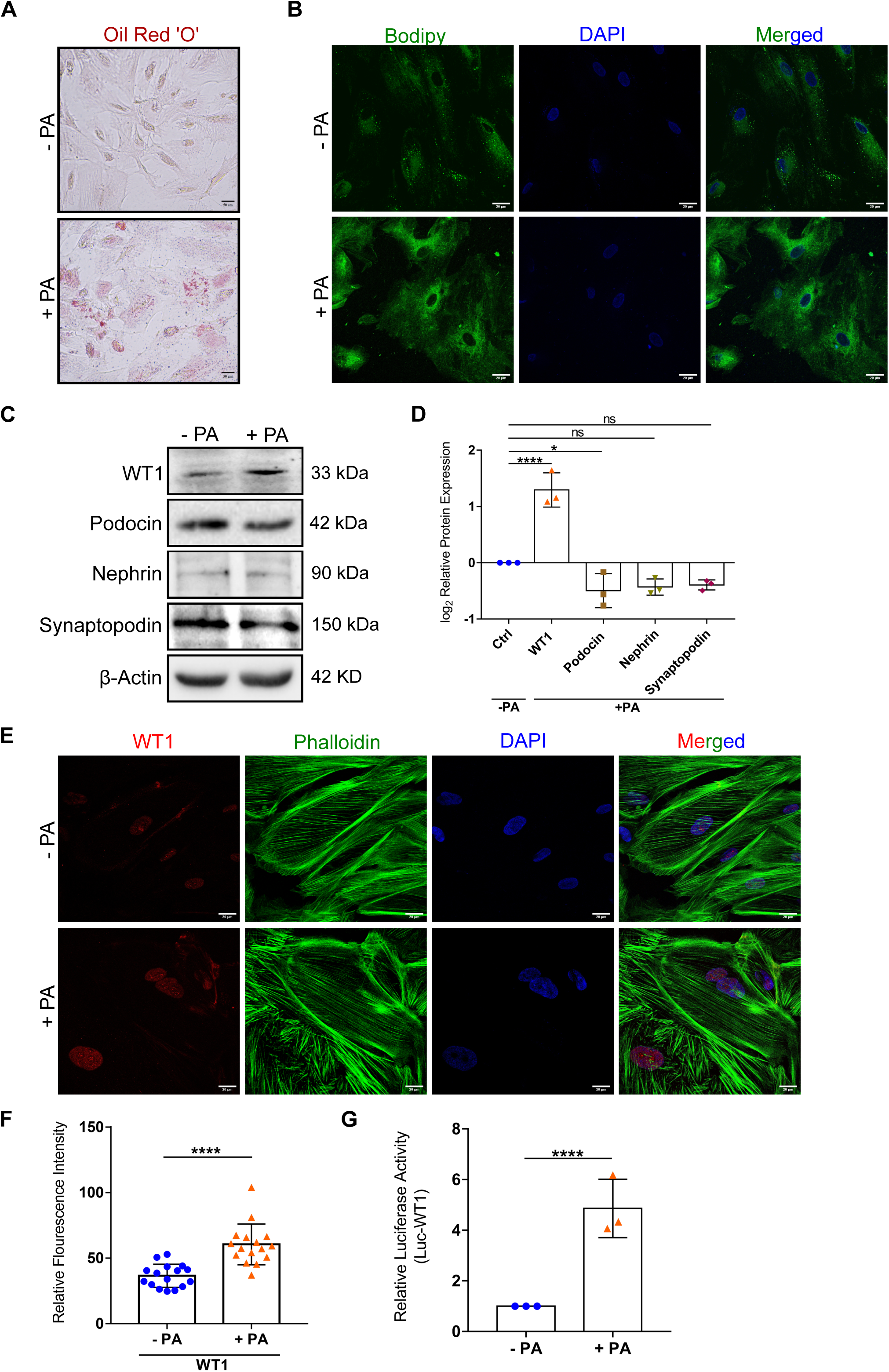
PA treated podocytes showed lipid accumulation and elevated WT1 expression: Immortalized human podocytes were treated with 100 µM of PA for 24 hours. **(A)** Oil Red ‘O’ staining of podocytes (Scale bar-5 µm). **(B)** Bodipy staining of podocytes (Scale bar-20 µm). **(C)** Immunoblotting analysis of podocytes lysate for WT1, podocin, nephrin and synaptopodin. **(D)**The immuno-blots were quantified using Image J from NIH and normalized to β-Actin. ****p<0.0001, *p=0.01 & ns=non significance (One- way ANOVA). **(E&F)** Immunoflourescence imaging of podocytes for WT1 (Scale bar-20 µm) and its measured relative fluorescence intensity using Image J from NIH ****p<0.0001 (unpaired t-test)**. (G)** WT1 promoter-Luciferase reporter assay ****p<0.0001 (unpaired t-test).

### PA treated podocytes showed elevated TGF-β1/Smad signaling

Podocytes are quiescent and are mono-nucleated. Since we observed binucleation in podocytes exposed to PA, we investigated the possible reasons. In an earlier study, it was shown that podocytes with elevated expression of endogenous TGF-β1 or exposed to TGF-β1 showed binucleation [12]. Interestingly, in the present study, we observed that PA treated podocytes showed elevated TGF-β1 expression and its downstream signaling molecules such as pSmad2 and 4 (Figure 2A&B). We also observed that the standard readouts of TGF-β1 expression; collagen and fibronectin were induced in PA treated podocytes (Figure 2A-D). WT1 exists in two major splice isoforms; “WT1 +KTS” and “WT1 −KTS” as alternative splicing at the end of exon 9 includes or skips three amino acids, lysine- threonine-serine (KTS). We observed TGF-β1 expression in podocytes that were ectopically expressing WT1 +KTS and WT1 −KTS, respectively (Figure 2E&F). The data suggests, elevated WT1 irrespective of its splice isoform is associated with slightly higher levels of TGF-β1 and it’s down-stream readouts such as α-SMA and fibronectin were elevated (Figure 2E&F).

**Figure 2:**
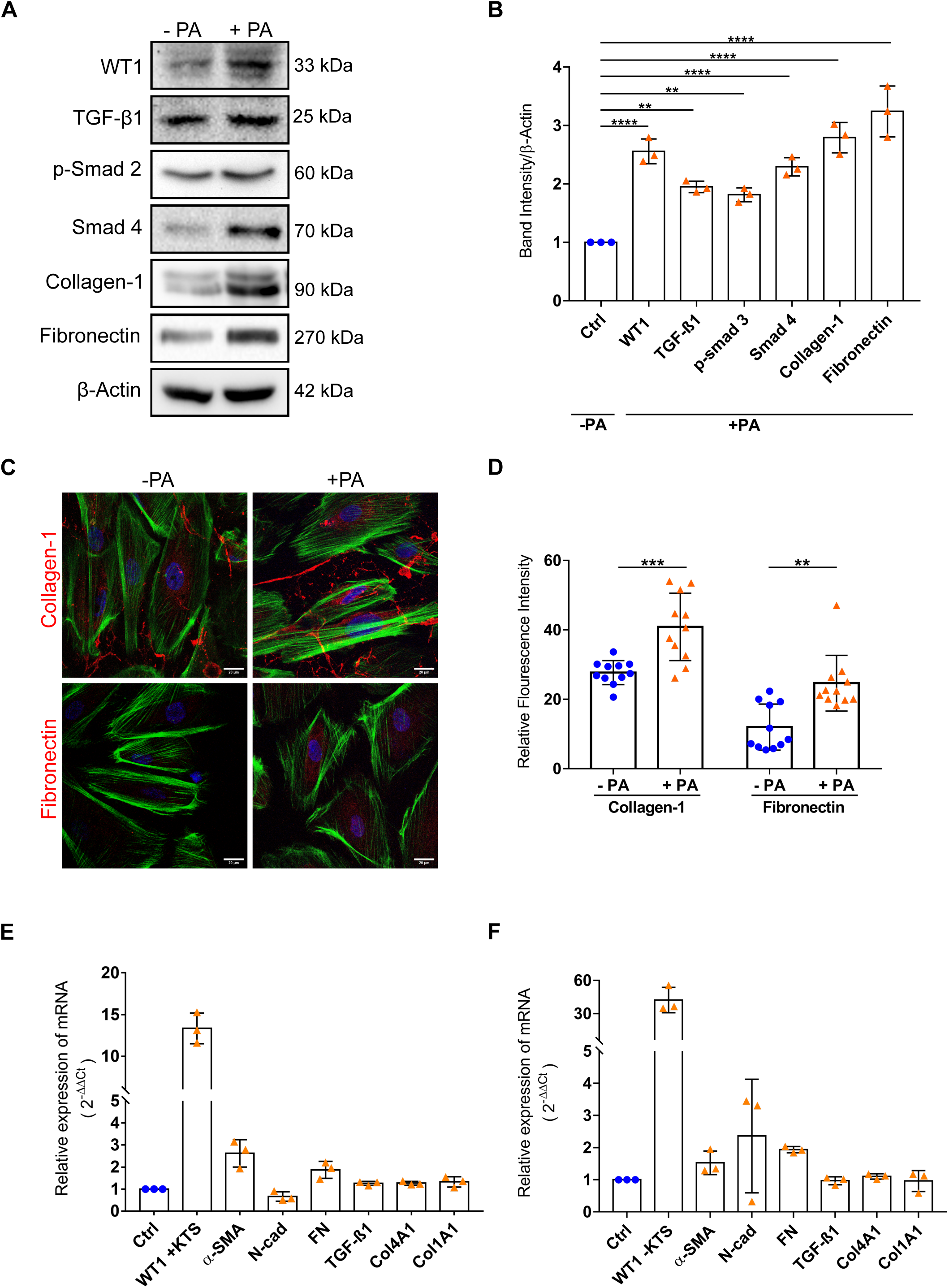
PA treated podocytes showed elevated TGF-β1/Smad signaling: **(A)** Immunoblotting analysis of podocytes lysate for WT1, TGF- β1, p-Smad 2, Smad 4, collagen-1 and fibronectin. **(B)** The immuno-blots were quantified using Image J from NIH and normalized to β-Actin. ****p<0.0001 **p=0.001 (One-way ANOVA). **(C&D)** Immunoflourescence imaging of podocytes for collagen and fibronectin (Scale bar-20 µm) and its measured relative fluorescence intensity using Image J from NIH ***p=0.0005 **p=0.001 (One-way ANOVA)**. (E&F)** Relative expression of mRNA using qRT PCR for ectopic expression of WT1 +KTS and –KTS isoforms. (One-way ANOVA).

### HFD fed mice showed elevated WT1 and TGF-β1 expression

In order to assess the effect of PA on glomerular biology and in turn kidney function, we fed the mice with 40% HFD diet for 16 weeks. These mice showed impaired kidney function as assessed by UCAR (Figure 3A) and subjecting urine to SDS-PAGE (Figure 3B). The data suggests impaired glomerular filtration in HFD fed mice. Immunoblotting and histochemical analysis revealed elevated WT1 and TGF-β1 in the glomerular preparations from HFD mice (Figure 3C-E). We also noticed elevated expression of TGF-β1 downstream markers *_a_*- SMA, fibronectin, and collagen (Figure 3C-E). The elevated expression of fibrotic markers is corroborated with glomerulosclerosis and degenerative changes as assessed by H&E, PAS, and Mason’s trichrome staining (Figure 3F). Together the data suggests elevated glomerular WT1 and TGF-β1 expression associated with compromised glomerular filtration. Since, TGF-β1 is a prosclerotic molecule, we observed sclerotic lesions in the HFD mice glomeruli in which elevated TGF-β1 expression was noticed.

**Figure 3:**
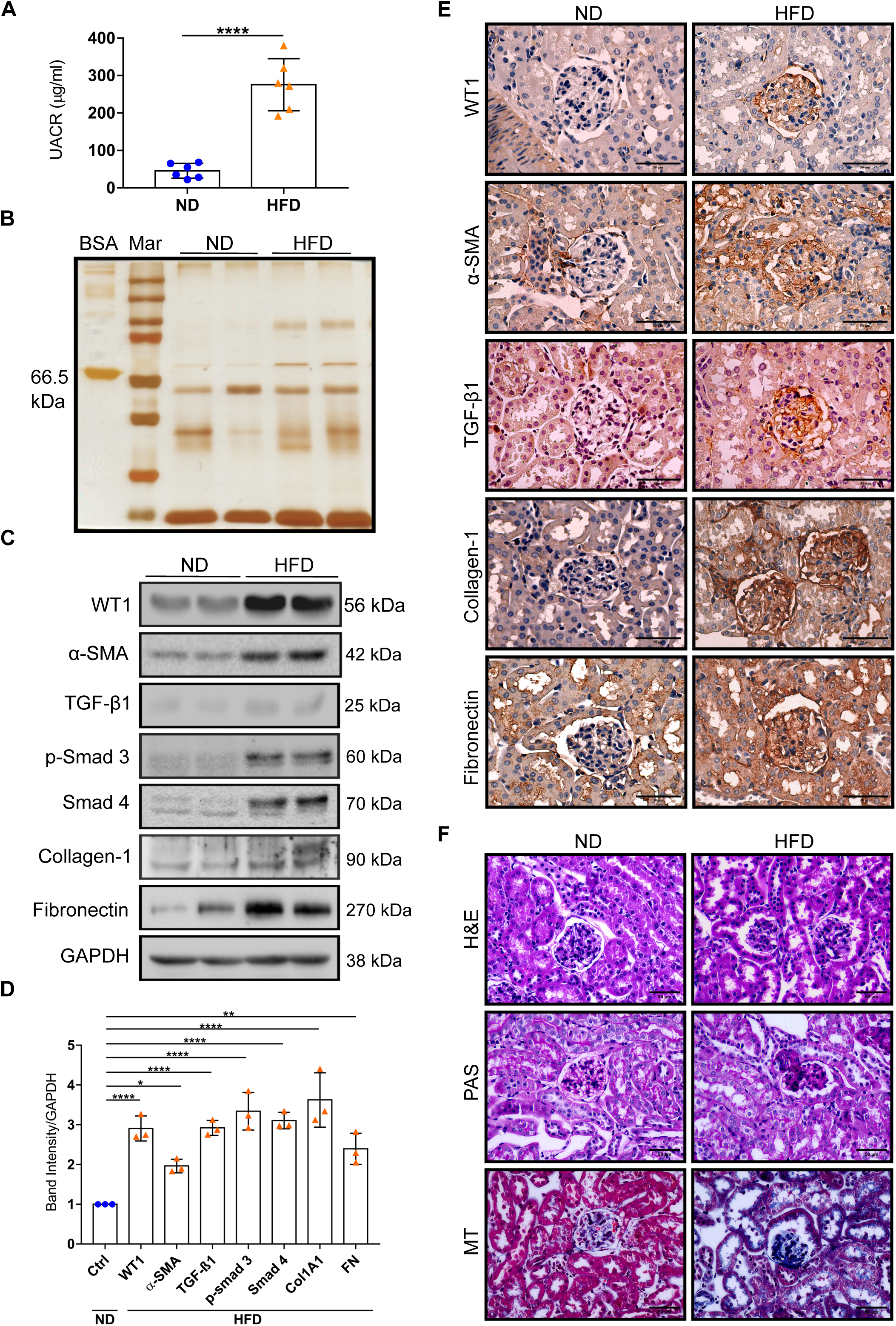
HFD fed mice showed elevated WT1 and TGF-β1 expression: **(A)** Urinary albumin creatinine ratio (UACR) of mice (n=6 for each group). ***p<0.0001 (unpaired t- test). **(B)** Urinary samples of age-matched ND and HFD mice were subjected to SDS- PAGE and Silver nitrate stained (n=2 for each group) **(C)** Immunoblotting analysis of podocyte enriched glomerular lysate for WT1, α-SMA, TGF- β1, p-Smad 3, Smad 4, collagen-1 and fibronectin. **(D)** The immuno-blots were quantified using Image J from NIH and normalized to GAPDH. ****p<0.0001 **p=0.001 *p=0.01 (One-way ANOVA). **(E)** Immunohistochemical staining of WT1, α-SMA, TGF- β1, collagen-1 and fibronectin (Scale bar-50 µm). Immunohistochemical analysis were performed on age-matched mice kidney sections, **(F)** H&E, PAS, and Masson’s trichrome staining. (Scale bar-50 µm).

### Elevated TGF-β1 expression evokes cell-cycle activation *in vitro* and *in vivo*

In an earlier study elevated growth hormone-dependent induction of TGF-β1 was associated with cell-cycle reactivation in podocytes [12]. Therefore, we next investigated the impact of PA-induced TGF-β1 expression on cyclins and cyclin-dependent kinases (CDKs) expression. We observed Cyc-B1 and Cyc-D1 expression in PA-treated podocytes in addition to CDK-4 expression (Figure 4A&B). Further, we noticed p53 and p21 expression in podocytes treated with PA. Since PA treated podocytes showed induction of cell-cycle, we next assessed the expression of cell-cycle activators in HFD fed mice glomerular preparation enriched with podocyte population. As anticipated, we noticed elevated expression of CDK4 and Cyc-B1 in podocyte lysate from the HFD fed mice (Figure 4C-E). Immunofluorescent images for Cyc-B1 revealed that few podocytes were binucleated suggests that the terminally differentiated cells are showing maladaptive response. Since podocytes considered to be quiescent and presented with constitutive expression of cell cycle inhibitors and minimal expression of cell cycle activators. Earlier studies revealed that presence of binucleated podocytes suggests cell cycle activation, nevertheless, these podocytes eventually undergo mitosis-associated cell death [16]. Glomerulus of HFD fed mice shows reduced count of podocytes suggests that cells may enter into catastrophe (Figure 4F).

**Figure 4:**
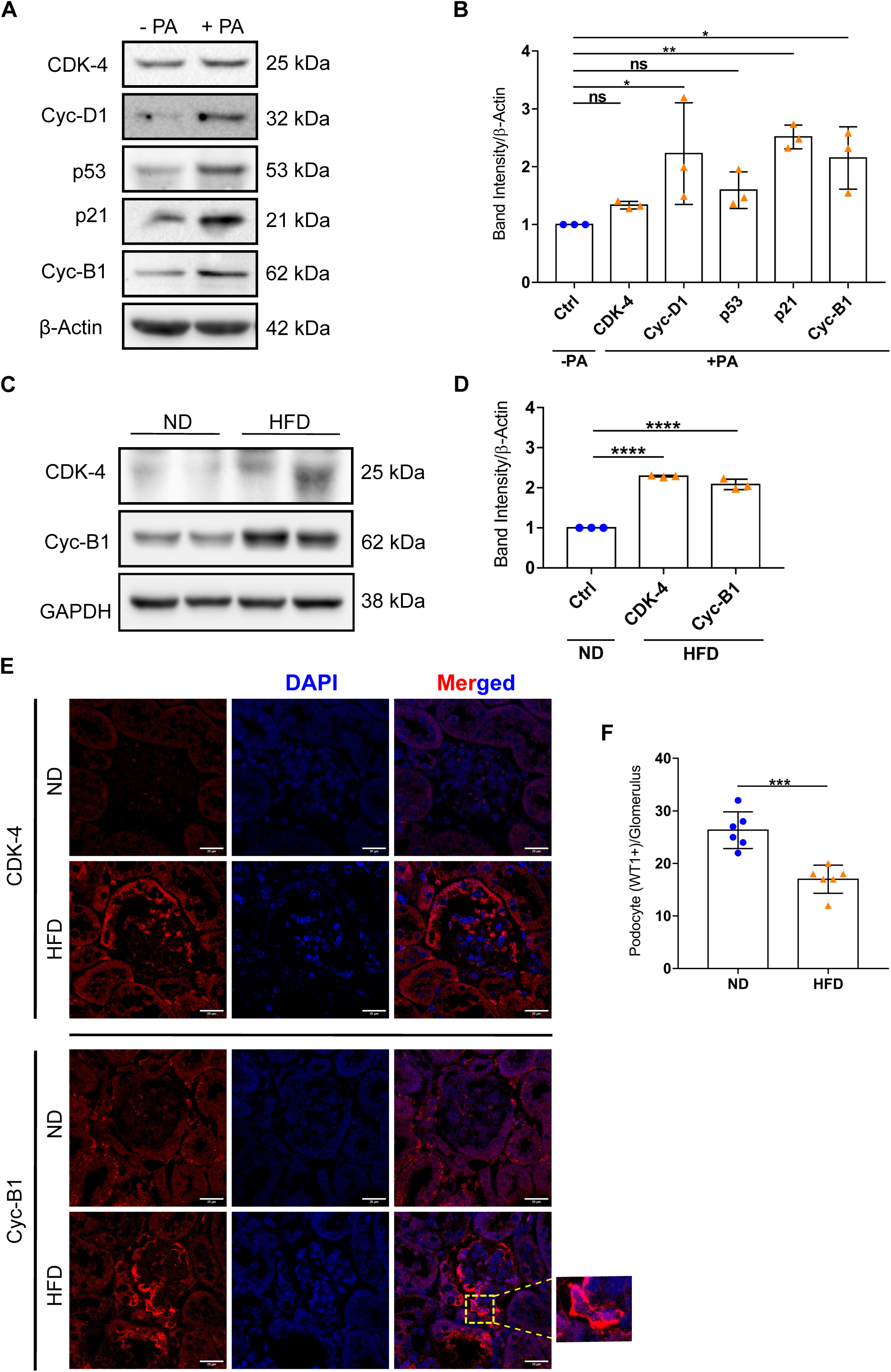
Elevated TGF-β1 expression evokes cell-cycle activation *in vitro* and *in vivo*: **(A)** Immunoblotting analysis of podocyte lysate for cell cycle markers. **(B)** The immuno-blots were quantified using Image J from NIH and normalized to β-Actin. **p=0.01 *p=0.01 p=ns (One-way ANOVA). **(C)** Immunoblotting analysis of podocyte enriched glomerular lysate CDK4 and Cyc-B1. **(D)**The immuno-blots were quantified using Image J from NIH and normalized to GAPDH. ****p<0.0001 (One-way ANOVA). **(E)** Immunoflourescence imaging of kidney tissues for CDK4 and Cyc-B1 (Scale bar-20 µm) **(F)** Podocyte count in each glomerulus ***p=0.0001 (unpaired t-test).

## Discussion

Our study revealed that exposure of human podocytes to PA resulted in the accumulation of lipid droplets. Lipid accumulation in podocytes was concomitant with elevated WT1 expression, as evidenced by both increased expression and increased promoter activity. Despite this increase in WT1 expression, PA-treated podocytes exhibited loss of podocyte-specific structural and cytoskeletal proteins, including nephrin, podocin, and synaptopodin, accompanied by cytoskeletal disruption. PA-exposed podocytes also exhibited increased TGF-β1 expression, accompanied by Smad2/Smad4 activation and upregulation of downstream markers such as fibronectin and collagen, indicating PA- induced TGF-β1 signaling. Consistent with these *in vitro* findings, podocyte-enriched glomeruli from HFD-fed mice displayed a similar induction of TGF-β1/Smad signaling and its downstream target genes. Furthermore, these HFD-fed mice showed glomerulosclerosis and impaired kidney function. Alongside elevated WT1 and TGF-β1 levels, both *in vitro* and *in vivo* models showed activation of cell cycle markers including Cyclin D1, Cyclin B1, CDK4, and p53/p21. These data suggest that Cyclin D1/CDK4 and Cyclin B1 drive quiescent (G0) podocytes to re-enter the cell cycle, whereas p53/p21 act to halt further progression. In addition to the aberrant expression of cell-cycle activators, we observed binucleation in a substantial proportion (∼30%) of podocytes exposed to PA. However, we did not observe an increase in podocyte number; rather, podocyte number decreased in mice fed an HFD.

Earlier studies have shown that ORG is characterized by glomerulomegaly, podocyte hypertrophy, reduced podocyte density, and secondary FSGS [17] [18]. Hyperfiltration, insulin resistance, inflammation, and ectopic lipid deposition have been proposed as drivers [19] [20]. Our data showed that, in addition to these mechanisms, particularly PA alone, could be sufficient to induce podocyte lipid droplet accumulation, loss of structural proteins, activation of TGF-β1 signaling, and cell-cycle dysregulation, and interestingly these changes are mirrored in HFD-fed mice with glomerulosclerosis, fewer podocytes, and proteinuria. The concurrent rise of WT1 and TGF-β1 in PA-treated podocytes suggests that WT1 contributes to TGF-β1 induction under lipotoxic stress. This would be unexpected, given that WT1 has been reported to repress the TGF-β1 promoter by competing with Egr-1 at a GC-rich response element in reporter assays [20], [21]. However, WT1 can act as either an activator or a repressor depending on cellular context [23], [24] and its activity is shaped by isoform, cofactor availability, post-translational modification, and subcellular localization [24]. We speculate that lipotoxic stress shifts WT1 toward a transcriptionally permissive state at the TGF-β1 locus, either directly via promoter engagement or indirectly via altered Egr-1 activity or other WT1 targets. We will investigate the temporal association between WT1 and TGF-β1 induction in PA-treated podocytes. Parallel induction of both genes by PA-driven ER stress or oxidative stress, and compensatory WT1 upregulation, could be possible alternative explanations. The relationship is further complicated by evidence that TGF-β1 reduces WT1 expression in podocytes [25], which suggests that the sequence of events may differ from the established pathway.

Our findings suggest that aberrant cell-cycle re-entry may be an additional route to podocyte loss in lipotoxicity, although the relative contributions of apoptosis and mitotic catastrophe remain to be defined. Podocyte re-entry into the cell cycle and the associated cell-cycle disorders are increasingly recognized phenomena [26]. Although mature podocytes are normally quiescent, injury-induced cell-cycle re-entry has been documented in collapsing glomerulopathy, focal segmental glomerulosclerosis (FSGS), diabetic nephropathy, and various genetic models [27]. Re-entry restricted to G1 can lead to hypertrophy, whereas progression beyond G1 without successful cytokinesis can lead to binucleation, mitotic catastrophe, and detachment [12], [28], [29]. The concurrent induction of cyclin D1, CDK4, cyclin B1, p53, and p21 and the ∼30% binucleation we observed closely resemble the profilin1-deficient model [30] and are consistent with these abortive cell-cycle events. Unlike in collapsing glomerulopathy, where CDK inhibitors fall [31], [32] but p21 was increased in our model, as reported in cellular FSGS and Adriamycin nephropathy [33] where it may restrain cell-cycle progression. Our WT1 results also differ from the current understanding: WT1 is typically reduced in FSGS and in response to TGF-β1 [25], whereas we observed increased expression WT1 alongside loss of its canonical targets. Possible explanations include a compensatory response, altered WT1 activity despite higher abundance, or lipotoxic changes in WT1 cofactors; these possibilities remain to be investigated in detail. It is noteworthy that podocytes respond to noxious stimuli by inducing transcription factors such as Notch1 and ZEB2, whose expression is low at baseline in adult podocytes [34], [35], [36], [37], [38]. This study has limitations. WT1 induction and its relation to increased TGF-β1 signaling have not been fully resolved, and our data establish an association rather than a linear pathway. Palmitate also acts through multiple routes, including ER stress, oxidative stress, and inflammation, which we did not separate. Future work should test whether inhibiting TGF-β1 signaling, modulating WT1, or blocking CDK4 preserves podocyte number and structure, and should define the fate of cells that become binucleated (apoptosis, senescence, or detachment). In conclusion, our findings show that lipotoxic stress induces WT1 and TGF-β1, disrupts the podocyte phenotype, and drives an abortive cell-cycle re-entry that is associated with binucleation and podocyte loss. These findings identify the WT1–TGF-β1–cell-cycle axis as a potential contributor to obesity- related glomerulopathy and a candidate target for intervention.

## Acknowledgments

AKP acknowledge the funding from ICMR and DBT (Govt of India) to his laboratory. AKP also acknowledge the infrastructural facilities established by DST- FIST and DBT-BUILDER in the Department of Biochemistry and School of Life Sciences, University of Hyderabad.

## DISCLOSURES

The authors declare no conflict of interest.

## Notes

### Competing Interest Statement

The authors have declared no competing interest.

